# Genome-wide association meta-analysis of cocaine dependence: Shared genetics with comorbid conditions

**DOI:** 10.1101/374553

**Authors:** Judit Cabana-Domínguez, Anu Shivalikanjli, Noèlia Fernàndez-Castillo, Bru Cormand

**Author notes:** Equally contributed (Noèlia Fernàndez-Castillo, Bru Cormand). **CORRESPONDING AUTHORS:** Bru Cormand. Departament de Genètica, Microbiologia i Estadística, Facultat de Biologia, Universitat de Barcelona, Avinguda Diagonal 643, edifici Prevosti, 3ª planta, 08028, Barcelona, Catalonia, Spain. Noèlia Fernàndez Castillo. Departament de Genètica, Microbiologia i Estadística, Facultat de Biologia, Universitat de Barcelona, Avinguda Diagonal 643, edifici Prevosti, 3ª planta, 08028, Barcelona, Catalonia, Spain.

## Abstract

Cocaine dependence is a complex psychiatric disorder that is highly comorbid with other psychiatric traits. Twin and adoption studies suggest that genetic variants contribute substantially to cocaine dependence susceptibility, which has an estimated heritability of 65-79%. Here we performed a meta-analysis of genome-wide association studies of cocaine dependence using four datasets from the dbGaP repository (2,085 cases and 4,293 controls, all of them selected by their European ancestry). Although no genome-wide significant hits were found in the SNP-based analysis, the gene-based analysis identified *HIST1H2BD* as associated with cocaine-dependence (10% FDR). This gene is located in a region on chromosome 6 enriched in histone-related genes, previously associated with schizophrenia (SCZ). Furthermore, we performed LD Score regression analysis with comorbid conditions and found significant genetic correlations between cocaine dependence and SCZ, ADHD, major depressive disorder (MDD) and risk taking. We also found, through polygenic risk score analysis, that all tested phenotypes can significantly predict cocaine dependence status: SCZ (*R*^2^=2.28%; *P*=1.21e-26), ADHD (*R*^2^=1.39%; *P*=4.5e-17), risk taking (*R*^2^=0.60%; *P*=2.7e-08), MDD (*R*^2^=1.21%; *P*=4.35e-15), children’s aggressiveness (*R*^2^=0.3%; *P*=8.8e-05) and antisocial behavior (*R*^2^=1.33%; *P*=2.2e-16). To our knowledge, this is the largest reported cocaine dependence GWAS meta-analysis in European-ancestry individuals. We identified suggestive associations in regions that may be related to cocaine dependence and found evidence for shared genetic risk factors between cocaine dependence and several comorbid psychiatric traits. However, the sample size is limited and further studies are needed to confirm these results.

## 1. Introduction

Cocaine is one of the most used illicit drugs worldwide and its abuse produces serious health problems. In Europe, around 5.2% of adults (from 15 to 64 years old) have tried cocaine (EMCDDA, 2017), but at most 20% will develop addiction (Wagner and Anthony, 2002). This information allows us to estimate the prevalence of cocaine dependence in the European population around 1.1%, similar to the prevalence observed in American populations (Compton et al., 2007).

Cocaine dependence is a complex psychiatric disorder that results from the interaction of environmental and genetic risk factors. It is one of the most heritable psychiatric conditions, with an estimated heritability of 65-79% (Ducci and Goldman, 2012). Although many case-control association studies in candidate genes have been performed, only a few risk variants for cocaine dependence have been identified and replicated so far, such as rs16969968 in the *CHRNA5* gene, encoding the cholinergic receptor nicotinic alpha 5 subunit, and rs806368 in *CNR1*, coding for the cannabinoid receptor 1 (Bühler et al., 2015). To date, only one genome-wide association study (GWAS) on cocaine dependence has been performed in European‐ and African-American individuals (Gelernter et al., 2014). When combining the two populations, one genome-wide finding was identified in the *FAM53B* gene, using a symptom count approach, but this hit could not be replicated in a subsequent study (Pineda-Cirera et al., 2018).

Several studies have shown that substance use disorders (SUD), and especially cocaine dependence, is highly comorbid with other psychiatric disorders and related phenotypes like aggressive, antisocial or risk-taking behaviors (Bezinović and Malatestinić, 2009; Hasin and Kilcoyne, 2012). For example, the occurrence of lifetime SUD in patients with schizophrenia (SCZ) is 70-80%, in attention deficit/hyperactivity disorder (ADHD) it is 39.2% and in major depressive disorder (MDD) it is 16.1% (Currie et al., 2005; Piñeiro-Dieguez et al., 2016; Westermeyer, 2006). Conversely, about 81% of substance abuse/dependence patients have at least one comorbid mental disorder: 33% MDD, 11% SCZ and 9% personality disorders (Shantna et al., 2009). Such comorbidity is associated with an increase of severity for all disorders, although it is unclear whether this relationship is causal or the result of shared genetic and/or environmental risk factors. Some studies have started to inspect these relationships using both genetic correlation and polygenic risk score approaches, supporting the hypothesized role of shared genetic risk factors in the lifetime co-occurrence of several psychiatric disorders and SUD (Carey et al., 2016; Du Rietz et al., 2017; Hartz et al., 2017; Reginsson et al., 2018).

Here we performed a GWAS meta-analysis of cocaine dependence in samples with European ancestry using datasets from the dbGaP repository. Then we investigated the shared genetics between cocaine dependence and other psychiatric traits.

## 2. Experimental procedures

Detailed description of the materials and methods used, as well as supplementary figures, can be found in the Supplementary Information.

### 2.1 Subjects

The case-control GWAS meta-analysis was performed using four datasets from the dbGaP repository (https://www.ncbi.nlm.nih.gov/gap) under the project 17170 (Table 1). All cases used met DSM-IV criteria for cocaine dependence, although most of them are also dependent to other drugs of abuse. Drug abuse or dependence were discarded only in controls of the SAGE sample, whereas in the other studies general population individuals were used as controls.

**Table 1.** Description of dbGaP samples used in the cocaine dependence GWAS meta-analysis

| Sample | N cases | %F cases | N controls | %F controls | Genotyping chip | dbGaP code |
| --- | --- | --- | --- | --- | --- | --- |
| Sample 1 (SAGE) | 468 | 39.3% | 1284 | 69.9% | Illumina ILMN_Human-1 | phs000092.v1.p1 (cases/controls) |
| Sample 2 | 609 | 45.9% | 410 | 50.2% | Illumina HumanOmni1-Quad_v1-0_B | phs000952.v1.p1 (cases/controls) + phs000179.v5.p2 (controls) |
| Sample 3 | 504 | 44.3% | 1190 | 62% | Illumina Human660W-Quad_v1_A | phs000277.v1.p1 (cases/controls) + phs000170.v2.p1 (controls) |
| Sample 4 | 504 | 36.7% | 1409 | 41.9% | Illumina HumanOmni1-Quad_v1-0_B | phs000425.v1.p1 (cases/controls) + phs000524.v1.p1 (controls) |
| **TOTAL** | **2,085** |  | **4,293** |  |  |  |

%F= percentage of females

Since samples 2-4 did not have enough controls to perform the association studies, we used controls from other datasets but genotyped with the same array and built using the same genome assembly. In all cases, patients and controls were from the same geographic area. In order to reduce bias, we merged controls with cases prior to quality control (QC) and imputation (Mitchell et al., 2014) (see Supplementary Information for detailed description of subjects used in this study).

The study was approved by the ethics committee of our institution, in accordance with the Helsinki Declaration and with the dbGaP protocols.

### 2.2 Quality control and association analyses

Prior to analysis, stringent QC was performed on the genotyped markers and individuals in each sample using the standardized pipeline “Ricopili” (Ripke, 2014). European-ancestry individuals were selected based on principal component analysis (PCA): PC1 and PC2 were used to define a genetic homogenous population, excluding individuals with PC values greater than three standard deviations from the reference population (European individuals from 1000 Genomes Project Phase 3 (1KGP3)). Related individuals and genetic outliers were excluded. A permutation test for between-group IBS differences with fixed 10,000 permutations was performed to discard population stratification between cases and controls.

After QC, non-genotyped markers were imputed using the European individuals from the 1KGP3 reference panel in MINIMAC3 (https://genome.sph.umich.edu/wiki/Minimac3).

For each sample, case-control GWAS was conducted using logistic regression under the additive model in PLINK v1.9 (http://pngu.mgh.harvard.edu/purcell/plink/). The 10 firsts PCs were included as covariates to correct for population stratification, and only variants with imputation INFO score >0.8 and minor allele frequency (MAF) >0.01 were considered.

### 2.3 GWAS meta-analysis

In total, 2,085 cases and 4,293 controls were meta-analyzed using an inverse-variance weighted fixed effects model in METAL (http://csg.sph.umich.edu//abecasis/Metal/)(Willer et al., 2010). Association results were considered only for variants with an effective sample size [N=2/((1/Ncases)+(1/Ncontrols))] greater than 70% of the full meta-analysis. Heterogeneity across studies was tested with the Cochran’s Q test and quantified with the I^2^ heterogeneity index, implemented in METAL.

Manhattan plot and Q-Q plot from each sample and the meta-analysis results were generated with the library “qqman” implemented in R.

### 2.4 LD Score intercept evaluation

LD score (LDSC) regression analysis was used to calculate LDSC intercept by regressing the chi-square statistics from GWAS against LD scores for each SNP (downloaded from GitHub website, https://github.com/bulik/ldsc)(Bulik-Sullivan et al., 2015b).

### 2.5 SNP heritability

LDSC 1.0.0 (https://github.com/bulik/ldsc/) was used to assess the proportion of phenotypic variance explained by common SNPs in the liability-scale, using a population prevalence for cocaine dependence of 1.1% (Compton et al., 2007).

Partitioned heritability analysis was performed based on 24 functional overlapping annotations described previously (Finucane et al., 2015). Enrichment in the heritability of a functional category was defined as the proportion of SNP heritability explained divided by the proportion of SNPs. The threshold for significance was calculated using the Bonferroni correction for multiple testing (*P*<2e-03).

### 2.6 Functional annotation of risk SNPs

SNPs were functionally annotated using FUMA (http://fuma.ctglab.nl/)(Watanabe et al., 2017). FUMA define lead SNPs as signals that are significantly associated with the phenotype of interest (we considered suggestive associations (*P*<1e-05)) and independent to each other (r^2^<0.1)). For each lead SNP, a “Genomic risk locus” is defined, including all independent signals that are physically close or overlapping in a single locus. The variants located in a “Genomic risk locus” were explored considering the following functional annotations: eQTL (from GTEx v6/v7 and BRAINEAC), CADD_v1.3, ANNOVAR, RegulomeDB_v1.1, 15-core chromatin state and GWAS-catalog e91.

### 2.7 Gene-based and gene-set association analyses

Gene-based and gene-set association analyses were performed with MAGMA 1.05b (Willer et al., 2010) using the summary statistics from the cocaine dependence GWAs meta-analysis. For gene-based analysis, the SNP-wise mean model was used as the statistic test, considering p-values for SNPs located within the transcribed region. For multiple testing corrections, 10% FDR was considered.

Gene-set analysis was used to test for enrichment in association signals in genes belonging to specific biological pathways or processes. We performed a competitive test using: “All Canonical Pathways” (1329 gene sets), “GO” (4436 gene sets) and “BioCarta” (217 gene sets) provided by MsigDB 5.1 (https://software.broadinstitute.org/gsea/msigdb/)(Subramanian et al., 2005). Multiple testing corrections were applied to each gene set separately. When gene sets are strongly overlapping, the Bonferroni correction can be quite conservative, and for that reason, we used an empirical multiple testing correction implemented in MAGMA, based on a permutation procedure.

### 2.8 Shared genetic factors between cocaine dependence and comorbid conditions

#### 2.8.1 Subjects

We studied shared genetic factors between cocaine dependence and six previously described comorbid conditions using publicly available GWAS summary statistics of SCZ, ADHD, MDD, children’s aggressive behavior, antisocial behavior and risk-taking behavior (Table 2). Summary statistics from the vitamin D levels GWAS of the UK Biobank was used as a negative control.

**Table 2.**
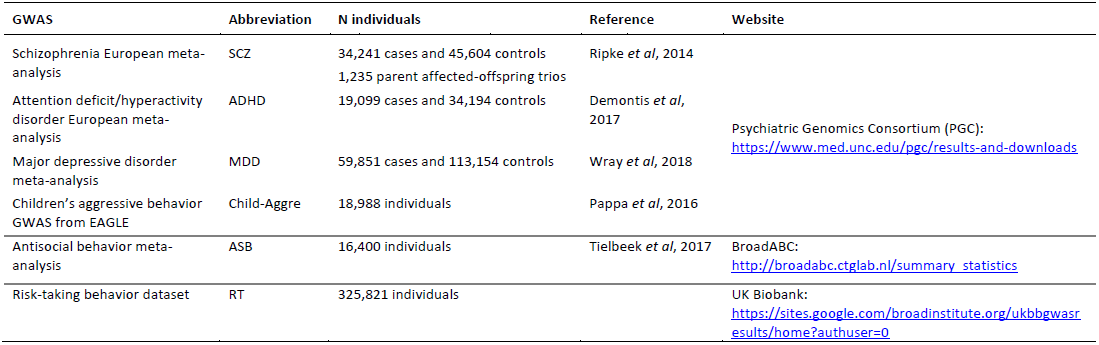
Description of samples used to inspect shared genetic factors between cocaine dependence and comorbid conditions

#### 2.8.2 LDSC Genetic correlation

Genetic correlation between cocaine dependence and the six selected comorbid disorders/traits was calculated using LDSC 1.0.0 (Bulik-Sullivan et al., 2015a). Summary statistics from all samples and pre-computed LD scores from HapMap3_SNPs calculated on 378 phased European-ancestry individuals from 1KGP3 were used (available at https://github.com/buligk/ldsc). After Bonferroni correction, the significance threshold was adjusted to *P*<7.1e-03.

Furthermore, the genetic correlation of cocaine dependence with other traits available at LD Hub (http://ldsc.broadinstitute.org/ldhub/)(Zheng et al., 2017) was evaluated (Bonferroni correction threshold *P*<7.2e-05).

#### 2.8.3 Polygenic risk scores for cocaine dependence

Poligenic risk score (PRS) analysis was performed using the PRSice 2.1.0 software (https://github.com/choishingwan/PRSice)(Euesden et al., 2015).

The four cocaine dependence datasets were merged using PLINK v1.9 and used as a target sample. After merging, strict QC was applied resulting in 5,957,307 SNPs in 2,083 cases and 4,287 controls. To assess population stratification PCA was performed, and the 10 first PCs and a dummy variable indicating genotyping-study was included in the PRS analysis as a covariate.

Summary statistics of the seven phenotypes described above were used as discovery samples, which were clumped (r^2^<0.1 in a 250-kb window) to remove SNPs in linkage disequilibrium (LD). Then, PRSs were estimated for each discovery sample using a wild range of meta-analysis p-value thresholds (P_T_) between P_T_=1e-04 and P_T_=1 at increments of 5e-05. For each P_T_, the proportion of variance explained (*R^2^*) by each discovery sample was computed by comparing the full model (PRS + covariates) score to a reduced model (covariates only). The reported *R^2^* value is the difference between *R^2^* from the two models. For quantitative traits we performed linear regression analysis, and for qualitative traits we used logistic regression and *Nagelkerke’s pseudo-R^2^* values are shown.

As recommended, we used the significance threshold of *P*=0.004 (Euesden et al., 2015). Bonferroni correction was applied considering the seven tested phenotypes (P<5.7e-04).

## 3. Results

### 3.1 GWAS results

We performed a GWAS meta-analysis of cocaine dependence using four datasets from the dbGaP repository. In total, we meta-analized 9,290,362 common genetic variants in 2,085 cases and 4,293 controls of European ancestry. No marker demonstrated significant heterogeneity between studies (Figure S1). The Q-Q plot (Figure 1A) displayed a λ of 1.06, comparable to other GWAS. The LDSC analyses estimated an intercept of 1.01 (SE=0.006; *P*=0.1), not significantly greater than 1, discarding residual population stratification or cryptic relatedness (Bulik-Sullivan et al., 2015b).

**Figure 1.**
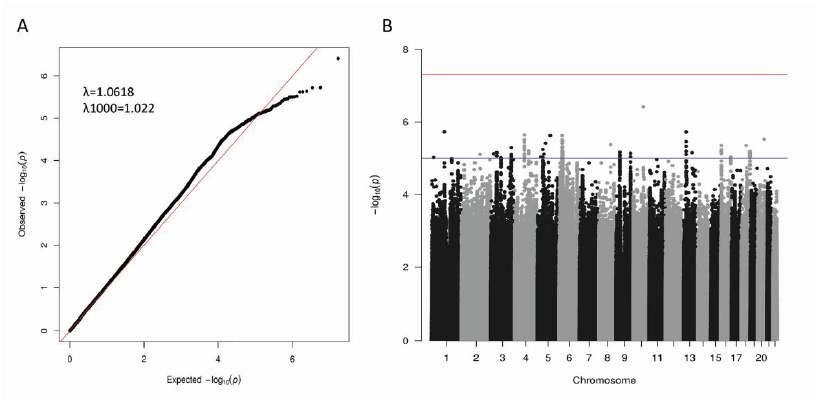
Results from the GWAS meta-analysis on cocaine dependence. A) Q-Q plot and B) Manhattan plot. Red line: threshold for genome-wide significance (*P* < 5e-08). Blue line: threshold for suggestive associations (*P* < 1e-05).

None of the analyzed variants reached the threshold for genome-wide significance (*P*<5e-08) in the SNP-based analysis, although we identified several suggestive associations (*P*<1e-05) (Table S1; Figure 1B).

### 3.2 Polygenic architecture of cocaine dependence

We applied LDSC analysis to assess the proportion of phenotypic variance explained by common SNPs. The estimated SNP heritability in liability-scale was h^2^_snp_=0.30 (SE=0.06; *P*=2.4e-07). Then we performed partitioned heritability analysis based on functional genomic categories and found significant enrichment in the heritability by SNPs located in intronic regions (enrichment=2.17; SE=0.45; *P*=1.2e-03), and a nominal result for conserved genomic regions (enrichment=23.63; SE=8.57; *P*=4e-03) (Figure S2).

### 3.3 Functional annotation of risk SNPs

To identify potentially interesting regions with FUMA we considered the SNPs showing suggestive associations (*P*<1e-05), as the SNP-based analysis did not reveal genome-wide significant hits (*P*<5e-08). We identified 23 lead SNPs which correspond to 22 genomic risk loci including 112 genes (Table 3). Interestingly, the risk locus located on chromosome 6 overlaps with a genomic region previously associated with schizophrenia. This region is defined by two lead SNPs (rs806973 and rs56401801, GWAs *P*=3.1e-06 and 3.4e-06, respectively) and it includes 77 genes and 458 nominally associated SNPs. Moreover, most of the SNPs in this region (447) are brain eQTLs for at least one member of a small group of 12 genes, including *BTN3A2*, *HIST1H2AK*, *ZSCAN31*, *PRSS16* and *ZNF184* (Figure 2 and S3).

**Figure 2.**
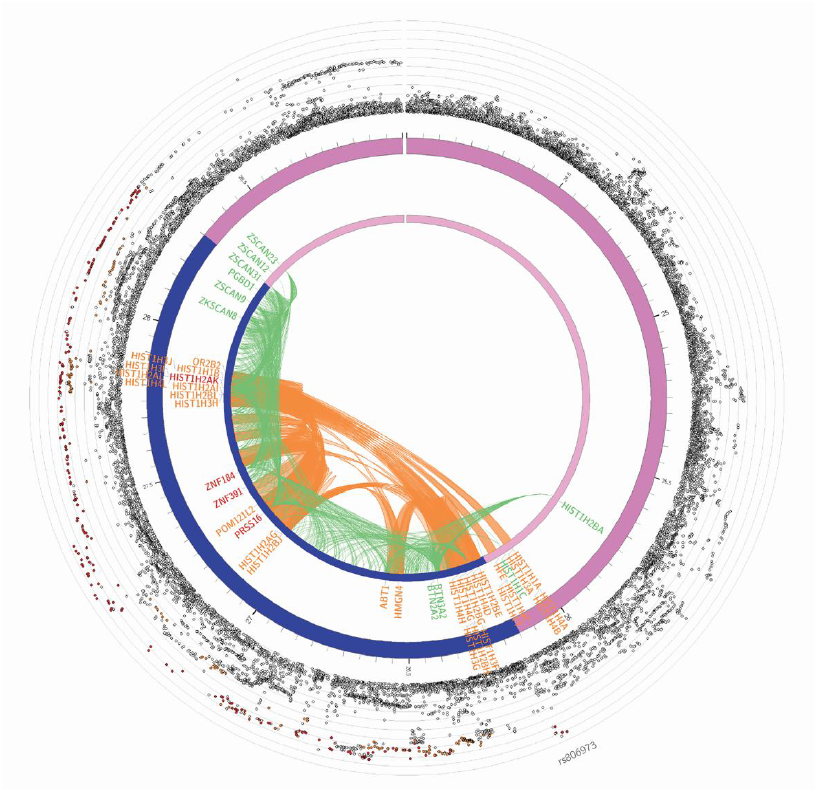
Circo-plot from genomic risk loci on chromosome 6 by FUMA. The most outer layer is the Manhattan plot (only SNPs with *P* < 0.05 are displayed). SNPs in genomic risk loci are color-coded as a function of their maximum r^2^ to the lead SNPs in the locus, as follows: red (r^2^ > 0.8), orange (r^2^ > 0.6), green (r^2^ > 0.4), blue (r^2^ > 0.2) and grey (r^2^ ≤ 0.2). The rs ID of the top SNPs in the risk locus is displayed in the most outer layer. Y-axis is ranged between 0 to the maximum ‐log_10_(p-value) of the SNPs. The second layer is the chromosome ring, with the genomic risk locus highlighted in blue. Here genes are mapped by chromatin interactions (orange) or eQTLs (green). When the gene is mapped by both, it is colored in red.

**Table 3.** FUMA analysis of genetic risk loci for cocaine dependence as identified from the GWAS meta-analysis

| Genomic Locus | Position* | Lead SNP | SNP position | GWAS P-value | nSNPs | nGenes | Genes |
| --- | --- | --- | --- | --- | --- | --- | --- |
| 1 | chr1:15916714-16049893 | rs148069235 | 15950064 | 9.52e-06 | 236 | 6 | *DNAJC16, AGMAT, DDI2, RSC1A1, PLEKHM2, SLC25A34* |
| 2 | chr1:104966872-104966872 | rs72685414 | 104966872 | 1.87e-06 | 1 | 0 |  |
| 3 | chr3:50615472-51356217 | rs148179194 | 50798652 | 7.03e-06 | 152 | 3 | *C3orf18, HEMK1, DOCK3* |
| 4 | chr3:84862871-84961810 | rs6767407 | 84955841 | 9.61e-06 | 27 | 0 |  |
| 5 | chr3:172170104-172214001 | rs57361543 | 172212144 | 5.09e-06 | 28 | 2 | *GHSR, TNFSF10* |
| 6 | chr4:82943149-83005137 | rs7675557 | 82970816 | 2.28e-06 | 16 | 1 | *RASGEF1B* |
| 7 | chr4:117560979-117607439 | rs67769911 | 117607070 | 6.33e-06 | 18 | 1 | *MIR1973* |
| 8 | chr5:30884915-31001774 | rs62357000 | 30884915 | 9.20e-06 | 4 | 1 | *CDH6* |
| 9 | chr5:44051305-44130776 | rs4410642 | 44129423 | 9.42e-06 | 61 | 0 |  |
| 10 | chr5:54436897-54754893 | rs334878 | 54519878 | 7.67e-06 | 90 | 9 | *ESM1, CDC20B, GPX8, MCIDAS, CCNO, DHX29, SKIV2L2, PLPP1, DDX4* |
| 11 | chr5:106788817-106796026 | rs71575441 | 106788817 | 2.37e-06 | 2 | 1 | *EFNA5* |
| 12 | chr6:26148326-28301195 | rs806973;<br>rs56401801 | 26148326;<br>27301762 | 3.14e-06;<br>3.44e-06 | 458 | 77 | Locus too broad |
| 13 | chr8:99193765-99226821 | rs4734387 | 99193765 | 4.20e-06 | 35 | 1 | *NIPAL2* |
| 14 | chr9:28890331-28993271 | rs35735220 | 28963289 | 6.86e-06 | 113 | 0 |  |
| 15 | chr9:118176789-118273407 | rs10121366 | 118244022 | 7.33e-06 | 54 | 0 |  |
| 16 | chr13:36947826-37019186 | rs79309473 | 36972202 | 1.89e-06 | 26 | 5 | *SOHLH2, SPG20, SPG20-AS1, CCNA1, SERTM1* |
| 17 | chr13:88150884-88150884 | rs7332726 | 88150884 | 7.10e-06 | 1 | 0 |  |
| 18 | chr16:6654017-6695032 | rs112252907 | 6675141 | 4.43e-06 | 42 | 1 | *RBFOX1* |
| 19 | chr16:84581684-84590225 | rs247831 | 84581893 | 9.17e-06 | 8 | 2 | *TLDC1, COTL1* |
| 20 | chr18:43206985-43231622 | rs1484873 | 43206985 | 4.45e-06 | 14 | 1 | *SLC14A2* |
| 21 | chr18:73743937-73775398 | rs73973283 | 73757906 | 6.46e-06 | 88 | 0 |  |
| 22 | chr20:54516338-54516338 | rs11086525 | 54516338 | 3.01e-06 | 1 | 0 |  |

\*Gene coordinates based on GRCh37/hg19. NSNPs: Number of nominally associated SNPs per genomic locus.

### 3.4 Gene-based and gene-set analyses

The gene-based analysis mapped 3,418,270 SNPs from the GWAS meta-analysis to 18,069 protein-coding genes (Figure S4 and Table S2). The *HIST1H2BD* gene, located in a genomic region on chromosome 6 that showed a suggestive association in the SNP-based analysis, displayed a significant gene-wise association with cocaine dependence (10% FDR), although it did not overcome the Bonferroni correction for multiple testing. Then we performed competitive gene-set tests for all BioCarta, GO and Canonical Pathways. No gene sets attained a significant association with cocaine dependence after correction for multiple testing (Table S3-5). However, interestingly, from the 10 GO gene sets with lower p-values, seven were related to synapse organization, glutamatergic neurotransmission and brain functions.

### 3.5 Cocaine dependence and shared genetic factors with comorbid conditions

Cocaine dependence is highly comorbid with other psychiatric disorders like SCZ, ADHD and MDD, and also with other phenotypes like aggressive behavior in children, antisocial behavior or risk taking. In order to investigate whether these phenotypic correlations are genetically mirrored, we performed a genetic correlation analysis using LDSC analysis and found significant genetic correlations (*P*<7.1e-03) between cocaine dependence and SCZ (rg=0.2; SE=0.05; *P*=1e-04), ADHD (rg=0.5; SE=0.08; *P*=1.6e-09), MDD (rg=0.4; SE=0.08; *P*=6.6e-07) and risk taking (rg=0.35; SE=0.06; *P*=9.1e-08) (Figure 3A). No significant correlations were found with children’s aggression or antisocial behavior, nor with a negative control (Vitamin D levels).

**Figure 3.**
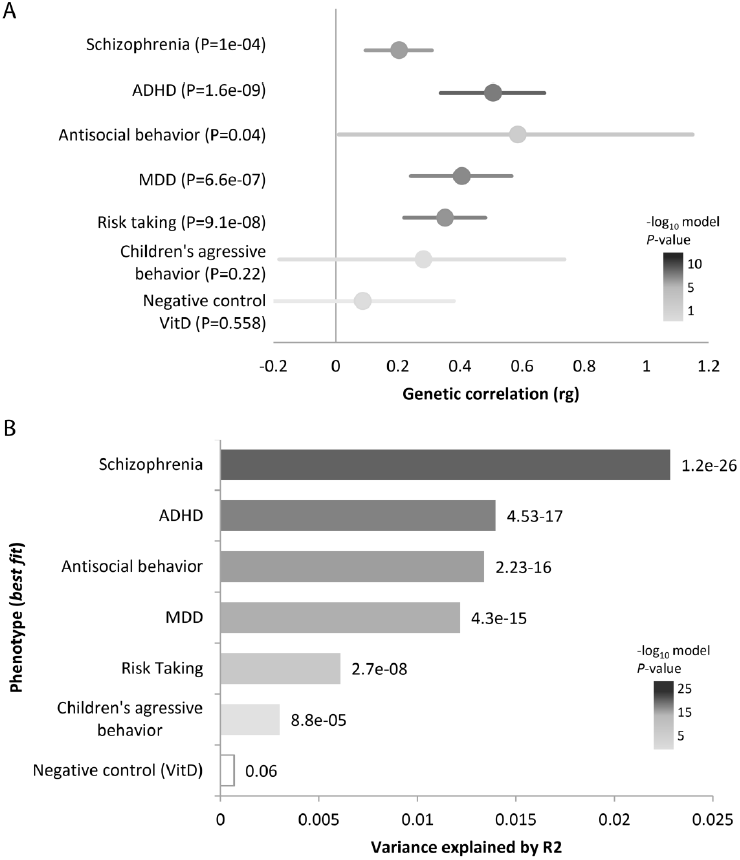
Shared genetic factors between cocaine dependence and comorbid conditions. A) Results from LD Score (LDSC) regression analysis showing genetic correlation (rg) between cocaine dependence and several traits. Error bars indicate 95% confidence limits. The significance threshold was set to *P* < 7.1e-03. B) Best fit results from Polygenic Risk Score (PRS) analysis for each tested phenotype. Values displayed next to each bar represent the p-value for significance for the most predictive models. The significance threshold was set at *P* < 5.7e-04.

On the other hand, we tested genetic correlations between cocaine dependence and all the GWAS summary statistics publicly available in the LD Hub. From the 690 tested traits, 109 demonstrated significant correlations after applying the Bonferroni correction, including a negative correlation with educational achievement (e.g. college completion) or with reproductive traits (e.g. age at first child) and a positive correlation with familiar situation (e.g. tobacco smoke exposure at home) or with several psychological or psychiatric traits like neuroticism, depressive symptoms or loneliness (Figure S5 and Table S6). The high number of significant associations may be explained by the high redundancy of the phenotypes of the LD Hub.

We also investigated the shared genetic etiology between cocaine dependence and comorbid phenotypes through PRS analysis, and tested whether these phenotypes can predict cocaine dependence status. For all the discovery samples tested, PRS significantly predicted cocaine dependence: SCZ (pseudo-*R*^2^=2.28%, P_T_=0.4911, *P*=1.21e-26), ADHD (pseudo-*R*^2^=1.39%, P_T_=0.04275, *P*=4.5e-17), antisocial behavior (pseudo-*R*^2^=1.33%, P_T_=0.4055, *P*=2.2e-16), MDD (pseudo-*R*^2^=1.21%, P_T_=0.0129, *P*=4.35e-15), risk taking (*R*^2^=0.60%, P_T_=0.00135, *P*=2.7e-08) and children’s aggressive behavior (*R*^2^=0.30%, P_T_=0.3552, *P*=8.8e-05) (Figure 3B and S6). In all cases, the quantile plot shows the positive nature of this relationship as cocaine dependence increases with greater polygenic load at each discovery dataset (Figure S7). As expected, the negative control based on vitamin D levels did not predict cocaine dependence (*R*^2^=0.07%, P_T_=0.03075, *P*=0.06).

## 4. Discussion

To our knowledge, this is the largest GWAS meta-analysis on cocaine dependence performed so far in individuals with European ancestry, although the sample size is still limited. No genome-wide significant (GWS) signals emerged from the SNP-based analysis, but the gene-based study identified *HIST1H2BD* as significantly associated with cocaine dependence. This gene is located in a region of chromosome 6 enriched in immune system and histone-related genes. These pathways have previously been associated with other psychiatric disorders like schizophrenia, major depressive disorder and bipolar disorder (O’Dushlaine, 2015). Despite the lack of GWS findings in this region, we identified many subthreshold variants. Based on the omnigenic model that has recently been proposed for complex disorders, these variants could point at regulatory elements of core genes (Boyle et al., 2017; Wray et al., 2018) and, therefore, contribute to the susceptibility to cocaine dependence, as most of them are brain eQTLs. Interestingly, this region overlaps with the genomic region that has been most consistently associated with schizophrenia. Indeed, it contains five SNPs (rs16897515, rs17693963, rs34706883, rs41266839 and rs55834529) nominally associated with cocaine dependence (*P*<1e-04) that had previously been associated with schizophrenia and with bipolar disorder, being the risk allele the same in all studies. The genetic variant most consistently associated with schizophrenia is rs17693963, reported in five different studies (Bergen et al., 2012; Ripke et al., 2011, 2014; Ruderfer et al., 2014; Sleiman et al., 2013), which is a brain eQTL for *PRSS16*, *ZSCAN9, ZNF184* and *ZSCAN31*. Furthermore, a transcriptomic study performed in lymphoblastoid cell lines of 413 schizophrenic cases and 446 controls found that the top differentially expressed genes are located in this region (e.g. *HIST1H2BD*, *HIST1H2BC*, *HIST1H2BH*, *HIST1H2BG* and *HIST1H4K*) (Sanders et al., 2013).

Cocaine dependence is a highly heritable disorder (around 65-79% (Ducci and Goldman, 2012)). The results from LDSC analysis showed that common SNPs significantly capture almost a half of the cocaine dependence heritability (h^2^_SNP_=0.3; *P*=2.4e-07). Similar results were obtained previously for alcohol dependence (Mbarek et al., 2015) and other psychiatric disorders (Lee, 2013). However, a recent study estimated a lower SNP-heritability for alcohol dependence (h^2^_SNP_=0.09 for European-American subjects) using a larger GWAS meta-analysis on alcohol dependence (Walters et al., 2018).

It is well known that most psychiatric disorders are highly comorbid. About 73.4% of cocaine abuse/dependence patients have comorbid mental disorders: 49.7% have personality disorders (e.g. 5.3% schizoid and 17% antisocial personality) and 61.5% other mental disorders (e.g. 23.4% MDD and 20.5% anxiety) (Arias et al., 2013). However, the reasons for these covariations remain largely unknown. We investigated whether the phenotypic correlations between cocaine dependence and six comorbid psychiatric traits are genetically mirrored by performing genetic correlation analyses using LDSC. For the first time we found significant genetic correlation with ADHD, SCZ, MDD and risk-taking behavior, although these results should be taken with caution and need to be followed-up in a larger sample of cocaine-dependent individuals. Furthermore, we used the PRS method that, in contrast to LDSC, uses individual-level SNP data, resulting in higher statistical power and allowing for direct testing of interaction effects. According to our results, all the tested comorbid conditions can predict cocaine dependence status, suggesting that cocaine dependence is more likely in individuals with many risk alleles for the tested conditions than in those with fewer risk alleles. To our knowledge, this is the first report of a shared genetic etiology between cocaine dependence and ADHD, antisocial behavior, risk-taking behavior and children’s aggressive behavior based on genome-wide data. Previous studies have reported significant PRS associations between cocaine dependence and SCZ or MDD (Carey et al., 2016; Hartz et al., 2017; Reginsson et al., 2018), and also between SUD and other psychiatric disorders (Du Rietz et al., 2017; Gurriarán et al., 2018), although our study used the largest sample of cocaine dependence for this type of analysis so far. This correlation can reflect biological pleiotropy, where similar genetic mechanisms influence more than one trait, or mediated pleiotropy, where one phenotype is causally related to a second phenotype, so that the variants associated with this phenotype are indirectly associated with the second one.

This study has strengths and limitations that need some discussion. We performed a GWAS meta-analysis using all the cocaine dependence datasets available at the dbGaP repository, but we could not find any GWS association at the SNP-based level, as expected given the limited sample size, with a total of around 6,000 subjects, one third of them cases. However, these data allowed us to detect genetic correlations between cocaine dependence and several co-occurring conditions. Also, we calculated polygenic risk scores that explain a small fraction of the variance in the target phenotype, with figures that are similar to those obtained for other pairs of psychiatric conditions. To obtain a more comprehensive picture of the etiological overlap between cocaine dependence and comorbid conditions, larger studies will be needed, and other genetic factors should be included in the analyses (e.g. CNVs and rare variants). On the other hand, some of the dbGaP datasets used included only cases but not control individuals. For this reason, we used controls from other datasets that can introduce potential biases into the experimental design. Nevertheless, we performed very strict quality controls to avoid population stratification: the paired case and control samples were genotyped with the same platform and come from the same geographical area, the merging of the different datasets was performed prior to quality control and imputation, and after that a permutation test was performed to discard population stratification (Mitchell et al., 2014). Moreover, the LDSC analyses confirmed that most of the observed inflation (λ =1.06) can be attributed to polygenicity rather than to residual population stratification or cryptic relatedness (Bulik-Sullivan et al., 2015b). Finally, the disease phenotype has not been excluded in most of the control samples, which may potentially dilute positive findings in the association study (but not lead to false positive results). However, based on the prevalence of cocaine dependence (about 1.1%), the probability of false negative results due to this effect is low, and similar controls were used in other GWAS of drug addiction (Ikeda et al., 2013; Johnson et al., 2016).

In conclusion, we reported the largest cocaine dependence GWAS meta-analysis on individuals of European ancestry, even though no GWS hits could be identified. Enlarging the sample size of this study would increase the chances to detect significant associations. However, the fact that our analyses highlighted a region on chromosome 6 that also pops-up in several schizophrenia GWAS supports the idea of shared genetic risk factors in these two comorbid disorders. This is in line with the significant results derived from the genetic correlation and PRS analyses in our study and in others. Finally, it would also be interesting to investigate the genetic pathways and neurobiological mechanisms that underlie the genetic overlap between cocaine dependence and comorbid traits.

## Author disclosures

### Contributors

JC-D and AS contributed to the acquisition of genotype data; JC-D performed the GWAS meta-analysis and secondary analyses with the assistance of AS; JC-D prepared the first draft of the manuscript and all figures and tables; BC and NF-C coordinated the study and supervised the manuscript preparation. All authors contributed to the design of the study and approved the final manuscript.

## Acknowledgments

We are really thankful to Paula Rovira, Marta Ribasés, Nina Roth Mota, Concepció Arenas and Oscar Lao for their support with the GWAS analysis. We are also very grateful to the developers of the software tools used, especially Stephan Ripke and Raymond Walters.

The authors acknowledge the contribution of data from dbGAP accessed through project number 17170:the Study of Addiction: Genetics and Environment (SAGE), one of the genome-wide association studies funded as part of the Gene Environment Association Studies (GENEVA) under GEI (supported by U01 HG004422, U01 HG004446, U01HG004438, HHSN268200782096C; and support for collection of datasets and samples was provided by the COGA; U10 AA008401, COGEND; P01 CA089392 and FSCD; R01 DA013423); GWAS of Heroin Dependence (supported by R01DA17305); GWAS of Cocaine Dependence in Two Populations (supported by RC2 DA028909, R01 DA12690, R01 DA12849, R01 DA18432, R01 AA11330, and R01 AA017535); CRIC Study (supported by U01DK060990, U01DK060984, U01DK061022, U01DK061021, U01DK061028, U01DK60980, U01DK060963 and U01DK060902); CIDR-Gelernter Study, a genome-wide association study funded as part of the Genetics of Alcohol Dependence in American Populations (supported by U01HG004438 and HHSN268200782096C); the COPDGene study (supported by U01HL089856 and U01HL089897, and the COPD Foundation through contributions made by an Industry Advisory Board comprised of Pfizer, AstraZeneca, Boehringer Ingelheim, Novartis, and Sunovion); and Personalized Medicine Research Project (PMRP) (supported by U01HG004608, U01HG004438 and U01HG004603).

